# TNFα and Endothelial IL-1 Receptor Signaling Drive Peritubular Capillary Regression and Fibrosis in Obstructive Kidney Injury

**DOI:** 10.64898/2025.12.30.697039

**Authors:** Charmain F. Johnson, Kate Wheeler, Jun Xie, Jenna Chan, George E. Davis, Courtney T. Griffin

## Abstract

Capillary regression destabilizes tissue homeostasis and contributes to chronic organ dysfunction, yet the inflammatory pathways that drive pathological vessel loss remain incompletely defined. We previously identified the inflammatory cytokines TNFα and IL-1 as conserved mediators of physiological vessel regression in the neonatal mouse eye, but whether these cytokines contribute to pathological capillary regression in adult mice is unknown. In this study, we investigated the capillary regression that occurs along with inflammation in murine kidneys following irreversible unilateral ureteral obstruction (UUO) surgical challenge. Mice lacking genes encoding global TNFα, the endothelial IL-1 receptor IL-1R1, or both (double knockout, DKO) were examined at 10 days after UUO surgery. While loss of the individual genes did not affect peritubular capillary (PTC) regression, PTC regression was significantly reduced in DKO mice. This reduction in PTC regression correlated with less expression of the tubular epithelial injury marker KIM-1. DKO kidneys also displayed less fibrosis by Picrosirius Red and Masson’s trichrome staining. These findings demonstrate that TNFα and endothelial IL-1R1 cooperatively drive pathological capillary regression in the irreversible UUO model of chronic kidney injury and that preservation of PTCs correlates with less renal tubular injury and fibrosis at 10 days after injury.

**NEW & NOTEWORTHY:** Pathological PTC regression drives progressive kidney injury, but its inflammatory triggers are unclear. Using the UUO model, this study identifies cooperative TNFα and endothelial IL-1R1 signaling as key drivers of capillary loss. While deletion of either pathway alone was insufficient, combined loss preserved PTCs, reduced tubular injury, and attenuated fibrosis. These findings highlight synergistic inflammatory signaling in microvascular loss and suggest dual targeting may help preserve PTC integrity in chronic kidney disease.

## INTRODUCTION

Capillaries play a central role in maintaining tissue and organ function by serving as the primary sites for the exchange of oxygen, nutrients, and metabolic waste products. This exchange is essential for sustaining tissue homeostasis and overall organ health. Capillary regression, which is also called rarefaction, disrupts this balance and contributes to organ dysfunction. Indeed, capillary loss is a hallmark of several pathological conditions, including diabetes mellitus, hypertension, and ischemic injury^1–3^. While pathological regression is commonly associated with disease, capillary regression also occurs as a tightly regulated physiological process. For example, soon after birth in mice, the ocular hyaloid vascular system undergoes coordinated regression while the retinal vasculature is established^4,5^. Similar physiological regression events are observed in other contexts of normal tissue remodeling, such as in the uterus during the menstrual cycle^6^. Our laboratories previously reported that TNFα and IL-1α/β are potent inducers of capillary regression in mouse hyaloids and in three-dimensional vascular networks generated from human umbilical vein endothelial cells (HUVECs) grown in collagen matrices^7^. However, whether these inflammatory cytokines also contribute to pathological capillary loss in adult mice remains unclear.

Chronic kidney disease (CKD) affects approximately 1 in 7 adults and is a major contributor to cardiovascular mortality^8^. Although CKD can arise from a variety of underlying conditions, loss of peritubular capillaries (PTCs) is among the earliest and most consistent structural abnormalities observed in both human biopsies and experimental models of kidney injury^9–11^. The unilateral ureteral obstruction (UUO) model is widely used to study progressive kidney fibrosis and exhibits robust interstitial inflammation, tubular injury, and PTC regression within the first week following obstruction^12^. TNFα and IL-1α/β are rapidly upregulated after UUO^13,14^, and these cytokines can negatively impact endothelial cell (EC) survival, promote oxidative stress, and suppress angiogenic signaling in other contexts^15–20^ raising the possibility that they may directly destabilize PTCs.

Based on our previous findings that TNFα and IL-1α/β promoted regression of HUVEC vascular networks and mouse hyaloids^7^, we hypothesized that these cytokines also contributed to pathological regression of PTCs during UUO. In addition, when we co-cultured HUVECs and pericytes, we found that death of HUVEC-lined tubes did not cause loss of associated pericytes^7^. In response to injury in the kidney, pericytes detach, migrate away from PTCs and differentiate into fibroblast or myofibroblasts and promote tissue fibrosis^9^. Therefore, we predicted that inhibition of endothelial signaling by TNFα and IL-1α/β in the UUO model could mitigate both capillary regression and subsequent renal fibrosis. To test this, we evaluated PTC density, tubular injury, and fibrosis in mice lacking global TNFα and endothelial-specific IL-1 receptor (IL1R1) signaling at 10 days after UUO surgeries. Our results indicate that TNFα and IL-1α/β signaling synergistically drive pathological PTC loss in the injured kidney. These findings reveal a novel role for inflammatory cytokines in destabilizing the renal microvasculature and highlight the pathological consequences of this process.

## MATERIALS AND METHODS

### Mouse lines and treatments

All mouse studies were conducted in accordance with the NIH *Guide for the Care and Use of Laboratory Animals* and approved by the Institutional Animal Care and Use Committee at the Oklahoma Medical Research Foundation (protocol #23-12). Mice were maintained on a C57BL/6J genetic background and housed in a single room within the animal facility under a 12-hour light/dark cycle, with food and water provided *ad libitum*. Mice used in this study include *Il1r1-flox* (No. 028398; The Jackson Laboratory)^21^, *Tnf^−/−^*(No. 003008; The Jackson Laboratory)^21^, *Cdh5(PAC)-Cre^ERT^*^2^ (gift of Ralf Adams, Max Planck Institute for Molecular Biomedicine; No. 13073; Taconic)^22^. *Tnf^+/−^* mice were crossed with *Tnf^+/−^* to generate *Tnf^+/+^*, *Tnf^+/−^*, and *Tnf^−/−^*littermates. *Il1r^fl/fl^* mice were crossed with *Cdh5(PAC)-Cre^ERT^*^2^ mice to generate *Il1r1^fl/fl^;iCdh5(PAC)-Cre^ERT^*^2^ mice (i.e., *Il1r1^iECKO^*), and Cre-negative *Il1r1^fl/fl^* littermates were used as controls (i.e., *Il1r1^WT^*). *Tnf^−/−^;Il1r1^fl/fl^*mice were crossed with *Cdh5(PAC)-Cre^ERT^*^2^ mice to generate *Tnf^−/−^;Il1r1^fl/f^;iCdh5(PAC)-Cre^ERT^*^2^ mice (i.e., DKO). Tamoxifen (Sigma, #T5648) was dissolved in peanut oil at 10 mg/mL and administered via intraperitoneal injection at 2 mg/day for 3 consecutive days to 8-week-old mice. Experiments were conducted 4 weeks after induction to allow for complete gene recombination. Note that *Tnf^+/+^*, *Tnf^+/−^*, and *Tnf^−/−^* littermates did not receive tamoxifen injections since they carried no inducible alleles. The sexes of mice used in each experiment are detailed in *Supplemental Table S3*. Genotyping was performed using primers listed in *Supplemental Table S1*.3

### Unilateral ureteral obstruction (UUO)

Irreversible UUO surgery was performed as previously described^23,24^. Briefly, 12-week-old mice were anesthetized with 3% isoflurane, placed on a heated surgical platform, and administered extended-release buprenorphine (1.0 mg/kg dose; ZooPharm; #1Z-74000-222510) as an analgesic. A mid-line 3 cm laparotomy was performed, and the left ureter was carefully isolated from surrounding tissues. The ureter was then ligated at two points using sterile absorbable suture (Ethicon; #VCP421 5-0) and transected between the ligatures to ensure complete and irreversible obstruction. The muscle and skin layers were closed using nylon nonabsorbable sutures (Monosof; SN-661G 5-0) and surgical wound clips (Roboz; RS-9262 9 mm), respectively. Animals were monitored daily for signs of distress. Mice were euthanized 10 days following UUO induction for kidney collection (both right and left kidneys) and analysis.

### Renal functional assays (serum BUN and creatinine)

Serum blood urea nitrogen (BUN) and creatinine were measured using dry-slide cartridges (BUN/CREA CLIP, IDEXX Laboratories). At 10 days following UUO, blood was collected by cardiac puncture, allowed to clot at room temperature for 20–30 min, and centrifuged at 2,000 × g for 10 min at 4 °C to isolate serum. Serum samples were diluted 1:4 by mixing 50 µL of serum with 150 µL of 1× Dulbecco’s phosphate-buffered saline (DPBS; ThermoFisher, #14190144) lacking calcium and magnesium, yielding a final volume of 200 µL per assay. BUN and creatinine concentrations were expressed in mg/dL.

### Immunofluorescence

Left and right kidneys were collected from euthanized mice, weighed, and fixed overnight in 4% paraformaldehyde at 4°C. Fixed tissues were cryoprotected by sequential incubation in 10%, 15%, and 20% sucrose solutions (1 hour each at 4°C), followed by overnight incubation in a 1:1 mixture of 20% sucrose and optimal cutting temperature (O.C.T.) compound (Sakura, #4583). The following morning, kidneys were embedded in O.C.T. using cryomolds and sectioned at 10 μm thickness using a cryostat (Epredia™ Microm HM525 NX Cryostat, Fisher Scientific).

Tissue sections were dried at 37°C, washed in PBS for 5 minutes to remove excess O.C.T., permeabilized in 0.3% Triton X-100 for 15 minutes, and blocked in 10% donkey serum (Jackson ImmunoResearch, #102644-006) in PBS for 1 hour at room temperature. Primary antibodies were diluted in 3% bovine serum albumin (BSA; Rockland, #BSA-50) and 0.1% Triton X-100 in PBS, and applied overnight at 4°C. The next day, sections were incubated with fluorophore-conjugated secondary antibodies diluted in 3% BSA in PBS for 1 hour at room temperature in the dark. Fluorescence images were acquired using a Nikon Eclipse Ti-E epifluorescence microscope, and image analysis was performed using FIJI (ImageJ) software. Primary antibodies used for immunofluorescence were goat-anti-KIM-1 (1:100, R&D Systems; #AF1817), rabbit-anti-cleaved caspase-3 (1:400, Cell Signaling; #9661), mouse-anti-αSMA-Cy3 (1:250; Sigma; #C6198) and goat-anti-CD31 (1:100, R&D Systems; #AF3628). Secondary antibodies used were Cy5-donkey-anti-rabbit IgG (1:500, Jackson ImmunoResearch), and Cy3-donkey-anti-goat IgG (1:500, Jackson ImmunoResearch). Hoechst (20 µg/mL) was added to the secondary antibody incubation.

PTC density was assessed as previously described^10^. Briefly, CD31-stained kidney sections were imaged (4 images per mouse), and a 15 × 15 grid (225 squares) was superimposed on each image. Squares containing CD31-positive staining were manually counted and expressed as a percentage of total grid squares (% of CD31⁺ grid). Tubular injury was assessed by measuring the area positive for the injury marker KIM-1. For each image, the total tubular area and KIM-1–positive area were quantified, and tubular injury was expressed as the percentage of KIM-1⁺ area relative to the total tubular area. For cleaved caspase-3 (CC3) and α-smooth muscle actin (αSMA), the total image area and the area positive for each marker were quantified from acquired images. Positive staining was expressed as the percentage of CC3⁺ or αSMA⁺ area relative to the total image area.

### Histological staining

Kidneys were fixed overnight in 4% paraformaldehyde at 4°C, dehydrated through a graded ethanol series, and embedded in paraffin. Sections (10 μm thick) were cut and mounted on glass slides for histological staining. Hematoxylin and eosin (H&E) and Masson’s trichrome staining were performed according to standard protocols to assess general tissue architecture and fibrosis, respectively. Brightfield images were acquired using a Nikon Eclipse 80i microscope equipped with a Nikon DS-Fi1 digital camera. Picrosirius Red (PSR) staining was performed using established protocols by the OMRF Imaging Core and polarized light images were acquired using a Zeiss Axioscan 7 digital slide scanner (Zeiss) located in the OMRF Imaging Core. The circular polarized light in the scanner was used to reliably acquire thick (mature) collagen fibers in red and thin (immature) collagen fibers in green^25,26^. Percent collagen area was calculated in eight regions of interest (ROIs) in polarized light images. To calculate relative content (red vs green) of total fibers, each image was converted from RGB to an Image5D format in ImageJ to allow separation of the red and green channels. For each polarized light image, six ROIs were selected, and the percent area occupied by red and green fibers was quantified. These values were averaged across ROIs and plotted as the relative content of total fibers.

### Statistical analyses

All statistical analyses were performed using GraphPad Prism (version 10.4.1). Data are presented as mean ± standard error of the mean (SEM), unless otherwise specified. Statistical significance was defined as *P* ≤ 0.05. Each dataset included at least *n* = 3 biological replicates, which was sufficient for inferential testing. For comparisons among more than two groups, one-way ANOVA with post-hoc testing was applied when a single independent variable was present and variances were equal (confirmed by F-test). For datasets with two independent variables and equal variances (confirmed by Spearman’s test), two-way ANOVA with post-hoc testing was used. When variances were unequal, data were log-transformed prior to two-way ANOVA.

## RESULTS

### Combined loss of TNFα and endothelial IL-1R1 signaling preserves peritubular capillaries following UUO

To determine the contribution of TNFα and IL-1R1 signaling toward peritubular capillary (PTC) regression, we subjected various genetic mouse strains to UUO surgeries. Mice that were analyzed included *Tnf^+/+^*, *Tnf^+/−^*, and *Tnf^−/−^* littermates generated from intercrosses of *Tnf^+/−^* animals. Additionally, *Il1r1^fl/fl^* (control = *Il1r1^WT^*) and *Il1r1^fl/fl^;Cdh5(PAC)-Cre^ERT^*^2^ (inducible EC *Il1r1* knockout = *Il1r1^iECko^*) littermates were generated for analysis of endothelial IL-1R1 signaling. Finally double knockout (*Tnf^−/−^;Il1r1^iECko^*= DKO) mice were generated for comparison against the control and single knockout genotypes listed above. 8-week-old mice were subjected to UUO surgeries. Note that *Il1r1^WT^* and *Il1r1^iECko^*littermates as well as DKO animals received tamoxifen (Tmx) 30 days prior to surgery to induce endothelial-specific deletion of *Il1r1*. UUO was performed on the left kidney (Figure 1A), and tissues were collected 10 days after surgery for analyses. As expected, the UUO kidney appeared visibly larger than the contralateral kidney from the same mouse (Figure 1A), consistent with hydronephrosis due to obstructive kidney injury.

**Figure 1:**
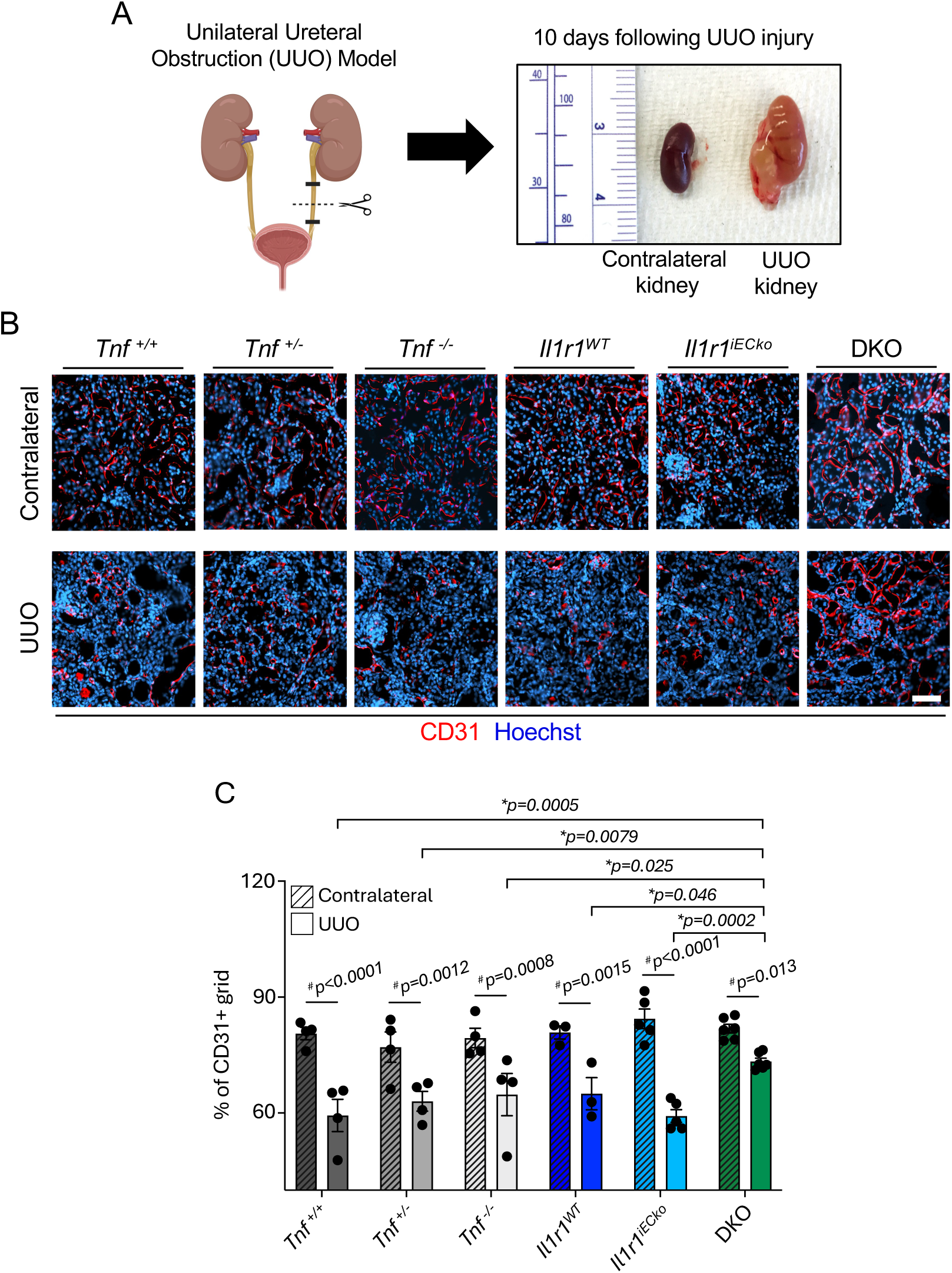
Synergistic TNFα and endothelial IL-1R1 signaling contribute to PTC regression following UUO. (A) Schematic of the unilateral ureteral obstruction (UUO) procedure and representative gross images of contralateral and hydronephrotic UUO kidneys from the same mouse collected 10 days after surgery. (B) Representative immunofluorescent images of PTCs stained for the endothelial marker CD31 (red) and counterstained with the nuclear dye Hoechst (blue) in contralateral and UUO kidneys across genotypes. Scale bar = 100 μm. (C) Quantification of CD31^+^ area in contralateral and UUO kidneys from the indicated genotypes. Each dot represents an individual animal; N=3-5. 3-4 images were quantified per animal. Data are presented as mean ± SEM. *p-values indicate significant differences between DKO UUO group versus UUO groups across other genotypes, # denotes p<0.05 between contralateral and UUO kidneys within each genotype. Data were analyzed using two-way ANOVA and post hoc comparisons. PTC=peritubular capillary; UUO=unilateral ureteral obstruction; DKO=double knockout (*Tnf^−/−^;Il1r1^fl/fl^;Cdh5(PAC)-Cre^ERT^*^2^).

We first measured kidney weights from contralateral and UUO kidneys across all genotypes used in this study. Although UUO kidneys weighed significantly more than contralateral kidneys, no differences were observed in the weights of UUO kidneys across the genotypes (Figure S1). Body weights remained relatively stable over 10 days post-UUO in all groups, with no significant differences observed between genotypes (Figure S2A). We also observed no differences in contralateral kidney to body weight ratios (Figure S2B), UUO kidney to body weight ratios (Figure S2C) and UUO to contralateral kidney weight ratios (Figure S2D) across genotypes. Next, to assess systemic kidney function following UUO, we measured serum blood urea nitrogen (BUN) and creatinine levels across genotypes. Neither BUN nor creatinine levels were elevated above the normal reference range (indicated by the two dotted lines) in any of the genotypes (Figure S3), which was predicted since the unmanipulated contralateral kidney can maintain normal kidney function in each mouse.

Quantification of PTC density using immunostaining for the endothelial marker CD31 (percent of CD31^+^ grid) revealed a significant reduction in PTC density in the cortex region of the UUO kidneys compared to contralateral kidneys across all genotypes (Figure 1C). We interpreted this as UUO-induced PTC regression based on published descriptions of the timing of this process^11^. When UUO kidneys were compared against each other, no differences in PTC regression were observed between *Tnf^−/−^* mice and littermate controls (*Tnf^+/+^* and *Tnf^+/−^*). Likewise, no differences were detected between *Il1r1^iECko^* and *Il1r1^WT^* littermate controls. However, combined loss of TNFα and endothelial IL-1R1 (DKO) resulted in significantly less PTC regression compared to all other groups, indicating protection against PTC regression following UUO challenge (Figure 1C). Representative images of immunostaining for the endothelial marker CD31, which was used to generate these quantitative findings, show denser and more continuous vascular networks in DKO UUO kidneys (Figure 1B).

### Expression of the proximal tubular injury marker KIM-1 is diminished with reduced capillary regression in DKO mice

PTCs are crucial for supporting proper tubular function by delivering oxygen and nutrients to renal epithelial cells. Damage or loss of PTCs can lead to local tissue hypoxia, which contributes to tubular injury and the progression of kidney disease^9,27^. To investigate the role of capillary regression in tubular injury following UUO, we assessed histological changes, cell death and tubular injury in the cortices of kidneys across genotypes. H&E-stained UUO kidney sections revealed tubular dilation (indicated by asterisks) in all groups (Figure 2A). To assess differences in cell death following UUO, we immunostained kidney sections for cleaved caspase-3 (CC3), a marker of apoptotic cells (Figure S4A). CC3^+^ cells were detected at low levels across all genotypes, and quantification of CC3^+^ areas revealed no statistically significant differences among the groups, although there was a downward expression trend in DKO kidneys (Figure S4B). Next, to quantify tubular injury, we analyzed UUO kidney sections for expression of Kidney Injury Molecule-1 (KIM-1), a biomarker of kidney tubular injury^28,29^, by immunofluorescence (Figure 2B). Although KIM-1 was broadly expressed in tubules across genotypes, the lowest KIM-1 staining was observed in DKO mice. Quantification of KIM-1 positive area normalized to total tubular area showed a significant reduction in the tubular injury marker in DKO mice compared to all other genotypes except the *Tnf^+/+^* group, which did not reach statistical significance (*p* = 0.060) (Figure 2C). Collectively, these findings suggest that PTC regression may contribute to tubular injury following UUO and that preserving PTCs could help mitigate tubular damage in kidney disease.

**Figure 2:**
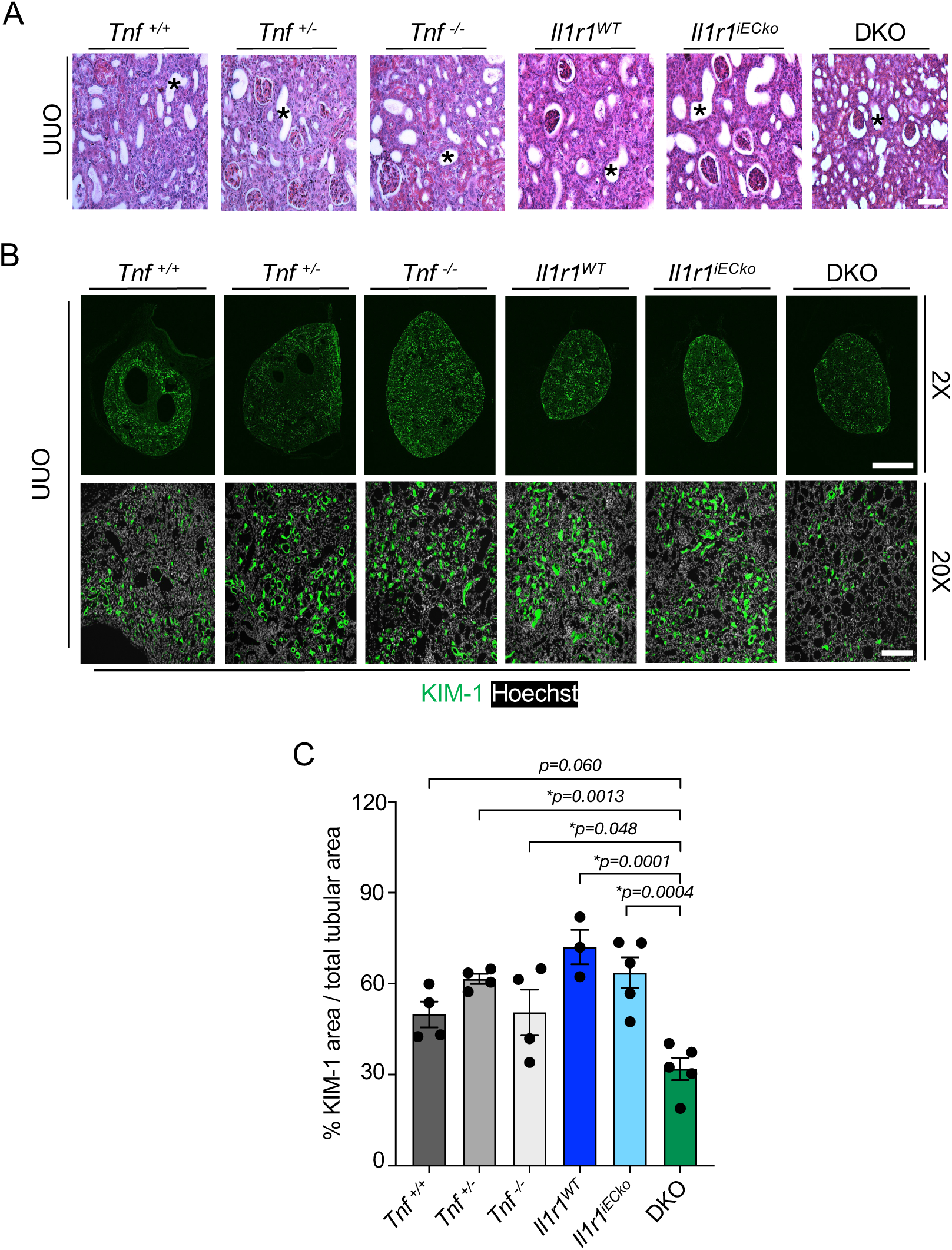
Proximal tubular injury is diminished with reduced capillary regression. (A) Representative H&E-stained kidney sections from the indicated genotypes 10 days after UUO. Asterisks (*) denote dilated tubules. Scale bar = 50 µm. (B) Representative immunofluorescence images of UUO kidneys stained for KIM-1 (green) and and counterstained with the nuclear dye Hoechst (white). Top row: low magnification (2X) whole-kidney images; scale bar = 1000 µm. Bottom row: high magnification images (20X); scale bar = 200 µm. (C) Quantification of KIM-1^+^ area normalized to total tubular area across genotypes. Each dot represents an individual animal; N=3-5. 3-4 images were quantified per animal. Data are presented as mean ± SEM. *p-values indicate significant differences between DKO UUO group versus other UUO groups. Data were analyzed using one-way ANOVA and post hoc comparisons.

### Reduced peritubular capillary regression correlates with less fibrosis

We previously speculated that capillary regression could drive subsequent tissue fibrosis^30^, which is prevalent in the UUO model and a significant contributor to kidney failure in CKD^31–33^.To determine whether reduced PTC regression alters the composition and organization of collagen fibers during fibrosis, we analyzed UUO kidney sections stained with Picrosirius Red (PSR) with circular polarized light microscopy. In contralateral kidneys across all genotypes, PSR staining revealed predominantly mature collagen fibers (appearing red; Figure 3A). UUO kidneys exhibited more PSR staining than contralateral kidneys, consistent with fibrotic remodeling (Figure 3B). Quantitative analysis of the percent collagen area in UUO kidneys was not different between DKO mice and all other groups (Figure 3C). However, quantitative analysis of mature (red) and immature (green) fibers in UUO kidneys demonstrated that double knockout (DKO) mice had significantly fewer mature collagen fibers and a corresponding increase in immature fibers compared with all other genotypes (Figure 3D). Additional histological staining using Masson’s trichrome revealed no obvious changes in collagen fiber content (blue staining) in contralateral kidneys between groups (Figure S5). However, when comparing UUO kidneys, those from DKO mice showed markedly less blue staining than those from all other groups (Figure S5). Unlike PSR-stained tissue sections imaged under circular polarized light, Masson’s trichrome predominantly stains mainly mature, thick collagen fibers, whereas thin collagen fibers are difficult to visualize^26^. This difference likely explains the reduced blue staining in UUO DKO kidneys compared with other UUO groups. Collectively, these findings highlight a correlation between PTC regression and renal fibrosis, which is attenuated in DKO mice.

**Figure 3:**
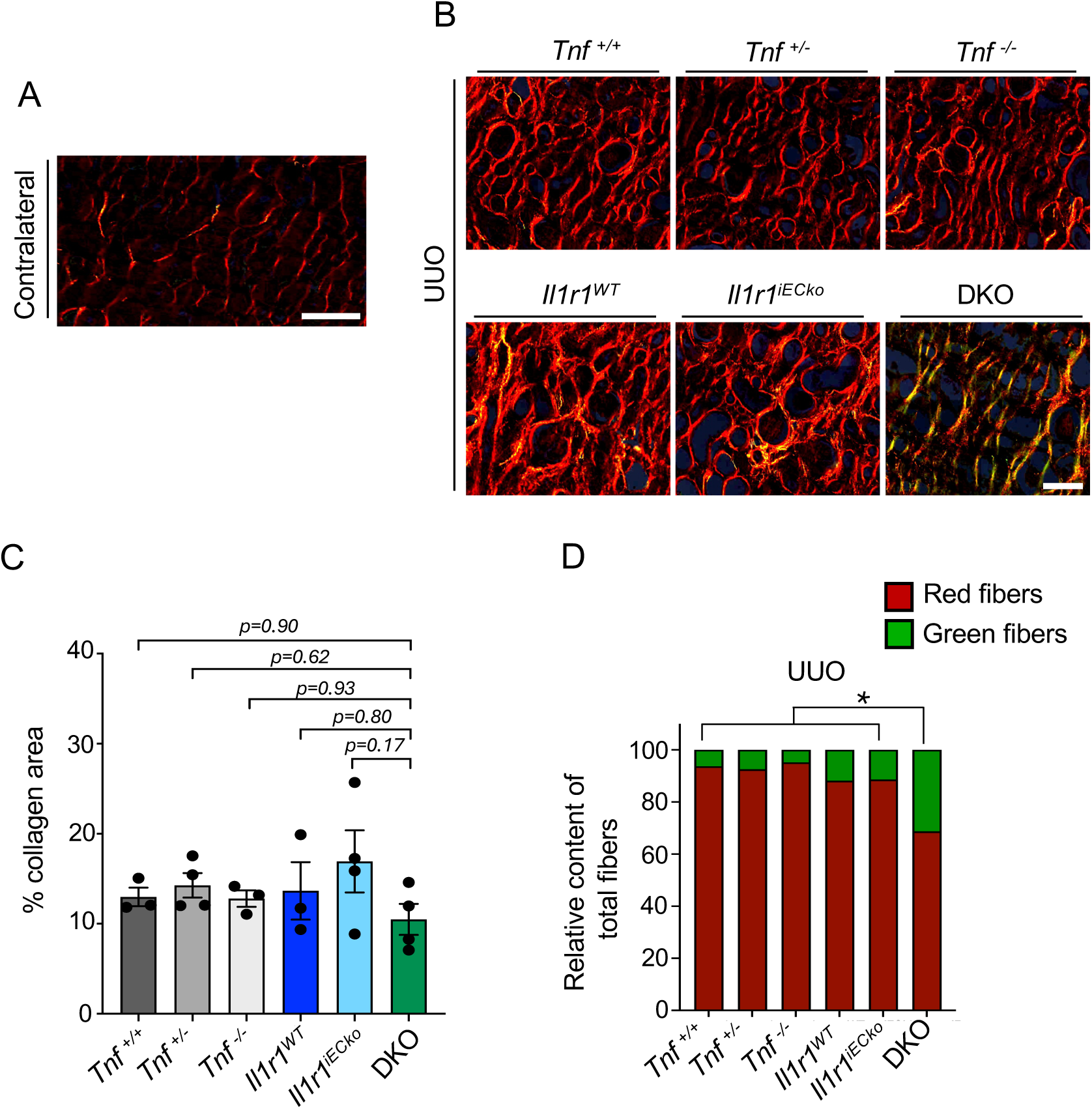
Loss of TNFα and endothelial IL-1R1 limits renal fibrosis. (A) Representative polarized light image of a contralateral kidney stained with Picrosirius Red. Scale bar = 50 µm. (B) Representative polarized light images of Picrosirius Red-stained UUO kidneys across genotypes. Scale bar = 50 µm. (C) Quantification of percent collagen area in UUO kidney across all genotypes. 8 ROI quantified per mouse. (D) Quantification of collagen fiber maturity in UUO kidneys from indicated genotypes. 6 ROI quantified per mouse. Mature (red) and immature (green) fibers are expressed as relative content of total fibers. Data are presented as mean ± SEM. *p-values indicate significant differences between DKO UUO group versus UUO groups across other genotypes. Data were analyzed using one-way ANOVA and post hoc comparisons

### The myofibroblast marker αSMA is elevated in DKO kidneys

Myofibroblasts are well-recognized for their synthesis of extracellular matrix (ECM) components and are major contributors to fibrosis of UUO kidneys^31^. We therefore questioned whether myofibroblast activation might explain the reduced fibrosis in DKO kidneys. Alpha-smooth muscle actin (αSMA) is a marker of myofibroblast activation that is upregulated after UUO challenge^34^, so we assessed its expression in contralateral and UUO kidneys in our different genetic models. In contralateral kidneys, αSMA immunostaining was minimal across all genotypes (Figure 4A). In contrast, UUO kidneys from all groups showed increased αSMA expression, with the most prominent staining observed in DKO mice (Figure 4A). Quantification of αSMA^+^ area confirmed that DKO UUO kidneys displayed significantly higher αSMA^+^ area compared to UUO kidneys from all other genotypes except the *Il1r1^iECko^* group, which did not reach statistical significance (*p* = 0.059) (Figure 4B). Therefore, DKO kidneys had more αSMA^+^ cells at 10 days after UUO injury despite displaying less fibrosis.

**Figure 4:**
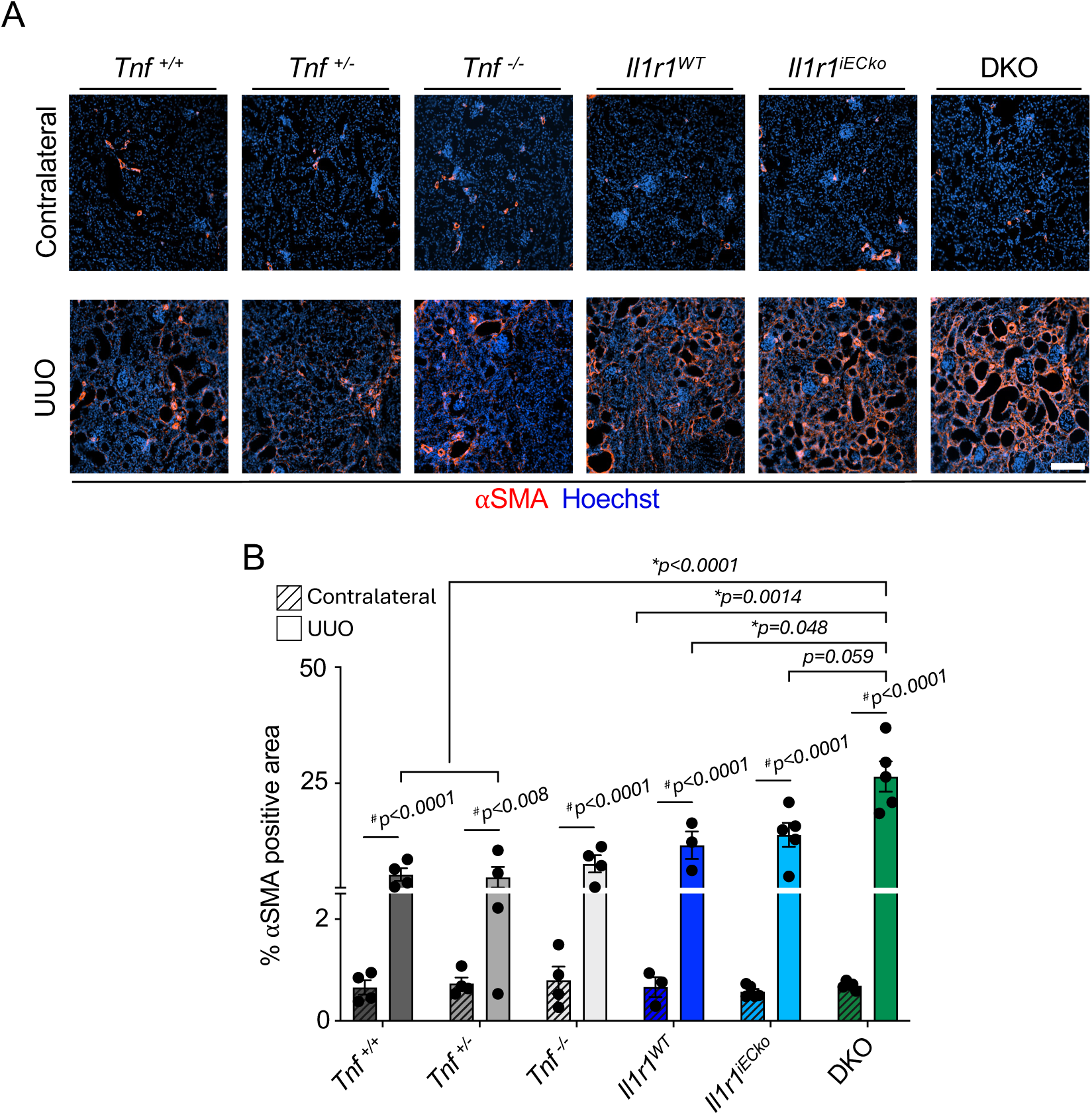
The myofibroblast marker αSMA is enhanced in mice lacking both TNFα and endothelial IL-1R1 signaling after UUO. (A) Representative immunofluorescence images of kidney sections stained for αSMA (red) and Hoechst (blue) from contralateral and UUO kidneys of indicated genotypes 10 days after UUO. Scale bar = 200 µm. (B) Quantification of αSMA-positive area (% of total area). Each dot represents an individual animal; N=3-5. 3-4 images were quantified per animal. Data are presented as mean ± SEM. *p-values indicate significant differences between DKO UUO group versus UUO groups across other genotypes, # denotes p<0.05 between contralateral and UUO kidneys within each genotype. Data were analyzed using two-way ANOVA and post hoc comparisons.

## DISCUSSION

In this study, we identify TNFα and endothelial IL-1R1 signaling as key drivers of pathological PTC regression following UUO injury in mice. While either pathway alone was dispensable, their combined loss significantly preserved PTC density following kidney injury. This vascular protection was accompanied by reduced tubular injury and diminished fibrosis, suggesting that PTC preservation mitigates downstream kidney damage. These findings extend our previous work implicating TNFα and IL-1 family cytokines in physiological vessel regression^7^ and establish their cooperative role in pathological capillary regression in obstructive kidney injury.

Loss of PTCs is an early and consistent feature of CKD and has been linked to chronic hypoxia, impaired nutrient delivery, tubular epithelial cell injury and progressive tubulointerstitial fibrosis^35–37^. By maintaining capillary density in the UUO model, DKO mice likely sustain better oxygen and nutrient exchange in the obstructed kidney, thereby reducing epithelial stress and injury, as reflected by lower KIM-1 expression. Similar relationships between vascular preservation and reduced tubular injury have been reported in UUO-challenged mice with genetic or viral overexpression of the pro-angiogenic factor angiopoietin-1 (*Angpt1*)^38–40^. This focus on angiogenesis-inducing agents stems from a recognition that UUO injury stimulates a burst of EC proliferation in PTCs that peaks at 4 days after injury^41^. However, EC damage also begins manifesting quickly after UUO injury, leading to PTC regression that progressively increases from 3-14 days after injury^11^. Therefore, EC proliferation and damage occur simultaneously in the UUO model, with cell death predominating by day 10. This is reminiscent of the differential pro-growth and pro-death EC effects attributed to TNFα and IL-1 in various studies and contexts^17,20,42–45^. Because we deleted *Tnf* and endothelial *Il1r1* one month prior to UUO injury in the present study, the increase in PTC density we detected at 10 days after injury indicates that these cytokines play a more consequential role in driving EC death than angiogenesis in the UUO model. We suspect that our failure to detect significant differences in EC apoptosis by CC3 immunostaining in control versus DKO kidneys at 10 days after UUO (Fig. S4B) likely means that we missed an earlier peak in cell death in this model.

In addition to PTC regression, the UUO model is also associated with substantial kidney fibrosis and has been used to define factors contributing to this damaging progress^23,32,46^. Although capillary regression and fibrosis both initiate soon after UUO injury, some reports indicate that regression exacerbates fibrosis in this model. For example, global knockout of the vascular development and maintenance factor *Angpt1* increases PTC regression and fibrosis in the UUO model^47^. Likewise, knockout of the urokinase plasminogen activator receptor (uPAR; *Plaur*), which facilitates ECM remodeling required for angiogenesis, yields adult mice with fewer PTCs, which experience enhanced capillary regression and fibrosis upon UUO challenge^48^. Correspondingly, there are several examples of mice in which both PTC regression and fibrosis are reduced after UUO injury. These include models overexpressing heme oxygenase-1^49^, genetically inhibiting signals that promote detachment of pericytes from ECs^41^, or receiving a pharmacological angiopoietin-2 inhibitor^38^. Our current study supports this growing body of evidence correlating PTC regression with kidney fibrosis by demonstrating less capillary loss and less fibrosis in DKO mice at 10 days after UUO injury.

Along with the decreased fibrosis we detected in DKO kidneys after UUO injury, we also observed an increased number of αSMA^+^ cells. This was unexpected because αSMA is a myofibroblast marker, and these synthetic cells are believed to be major contributors to ECM production and fibrosis in UUO-injured mice^31^. However, αSMA is also expressed by vascular smooth muscle cells and pericytes^50^. Given the preservation of PTCs in DKO mice, the elevated αSMA expression we detected may reflect increased mural cell coverage or contractile stabilization of the vasculature rather than an expansion of ECM-producing myofibroblasts. Consistent with this, EC-derived factors such as PDGF-B may promote mural cell recruitment under UUO conditions. Nevertheless, this still leaves the question of how to account for the decreased interstitial fibrosis in DKO kidneys. One possibility is that damaged ECs contribute directly to fibrotic matrix deposition. Ultrastructural studies have revealed that EC and PTC basement membrane thickening occurs soon after UUO injury^11^, suggesting a pro-synthetic endothelial phenotype. In support of this, mice with genetically reduced endothelial TGF-β signaling display significantly reduced tubulointerstitial fibrosis after UUO injury^51^. Because IL-1 and TNF signaling can synergize with TGF-β to promote an endothelial-to-mesenchymal transition^52,53^, future studies could investigate this process in UUO-challenged DKO mice. Notably, our previous study identified both TGFβ1 and TGFβ2 as pro-regressive factors for EC tube networks in vitro^7^, suggesting that endothelial TGFβ signaling may simultaneously contribute to kidney fibrosis and capillary loss under pro-inflammatory UUO conditions.

Our research naturally raises questions about the mechanisms through which TNFα and endothelial IL-1 signaling promote PTC regression in the UUO model. While follow-on work will be required to address these questions directly, our prior research may provide guidance for future studies. For example, our analysis of three-dimensional HUVEC networks undergoing TNFα- and/or IL-1β-mediated regression revealed signature signaling events, including increased expression of intercellular adhesion molecule-1 (ICAM-1), phospho-p38, and phosphorylated myosin light chain-2 (MLC2) and decreased expression of phospho-Pak2, acetylated tubulin, phospho-cofilin, and pro-caspase3^7^. Moreover, we described a combination of pharmacological agents that blocked TNFα/IL-1β-mediated regression in this HUVEC model^7^, which warrants testing in the UUO model. In a separate study, we discovered that TNFα drives rapid degradation of the pro-survival endothelial transcription factor ERG in mouse lungs^21^. This is interesting because we also found that endothelial ERG is downregulated in regressing murine and human ocular capillaries^54,55^, and its genetic overexpression in ECs can prevent retinal capillary regression in mouse models of retinopathy^55^. Therefore, future studies should address the possibility that endothelial TNFα/IL-1 signaling drive the degradation of pro-survival transcription factors like ERG and subsequent regression of PTCs in the UUO model. Such a pathway could provide multiple points of targeted intervention in the progression of CKD.

Altogether, this study demonstrates the contribution of TNFα and endothelial IL-1 signaling to pathological renal capillary regression. Because these inflammatory cytokines are upregulated in a wide variety of acute and chronic diseases associated with capillary regression, including diabetes, COVID-19, Alzheimer’s disease, heart failure, and sepsis^56–60^, our findings in the current UUO study may hint at their broader contribution to pathological capillary regression. We believe that studying the contribution of endothelial TNFα/IL-1 signaling to capillary regression in more disease models and organs will expand our understanding of the regression process and reveal new ways of intervening with the progression of ischemic tissue damage.

## LIMITATIONS OF STUDY

While we used an endothelial-specific model to delete the IL-1 receptor *Il1r1* in this study, we did not have access to a comparable model to delete TNFα receptors on ECs. Therefore, the phenotypes we detected in DKO mice may have been influenced by TNFα signaling in other cell types besides ECs. Another limitation of this study is that it did not address the contribution of thrombin signaling to UUO-induced PTC regression. This is relevant because our previous study with three-dimensional HUVEC networks demonstrated that thrombin enhances TNFα/IL-1β-mediated regression^7^. Notably, the thrombin inhibitor dabigatran etexilate significantly reduces fibrosis in UUO-injured mice^61^, and mice with global reduction of the thrombin receptor PAR4 likewise have less UUO-induced fibrosis^62^. Therefore, we believe investigating the combinatorial impact of endothelial TNFα/IL-1/thrombin signaling on PTC regression is a promising future direction. Finally, the single post-UUO timepoint at which we conducted analyses in the current study (day 10) only lends itself to correlative conclusions about the relationship between PTC regression and fibrosis or epithelial cell injury. A longitudinal model with analyses at early and late timepoints would facilitate more meaningful insights into the relationship between cytokine-induced PTC regression and its consequences for CKD progression.

## ACKNOWLEDGEMENTS

We thank all members of the Griffin laboratory for valuable feedback on this manuscript. Figure 1A and the graphical abstract were created using BioRender.com.

## SOURCES OF FUNDING

This work was funded by grants from the National Institutes of Health to C.T.G. (R35HL144605).

## DISCLOSURES

The authors declare no competing interests.

## AUTHOR CONTRIBUTIONS

C.F.J., G.E.D., and C.T.G. conceived and designed research. C.F.J., K.W., and J.C. analyzed data. C.F.J., K.W., and J.X. performed experiments. C.F.J., J.X., and C.T.G interpreted results of experiments. C.F.J. prepared figures and drafted the manuscript. C.F.J., K.W., G.E.D., and C.T.G. edited and revised the manuscript. C.F.J., K.W., J.X., J.C., G.E.D., and C.T.G. approved the final version.

## ABBREVIATIONS

αSMA: Alpha-Smooth Muscle Actin
CC3: Cleaved Caspase-3
CKD: Chronic Kidney Disease
EC: Endothelial Cell
IL-1: Interleukin-1
PSR: Picrosirius Red
PTC: Peritubular Capillary
TNF: Tumor Necrosis Factor
UUO: Unilateral Ureteral Obstruction

## Notes

### Competing Interest Statement

The authors have declared no competing interest.

## REFERENCES

1 Basile, D. P. Rarefaction of peritubular capillaries following ischemic acute renal failure: a potential factor predisposing to progressive nephropathy. Curr Opin Nephrol Hypertens 13, 1–7 (2004). 10.1097/00041552-200401000-00001

2 Battegay, E. J., de Miguel, L. S., Petrimpol, M. & Humar, R. Effects of anti-hypertensive drugs on vessel rarefaction. Curr Opin Pharmacol 7, 151–157 (2007). 10.1016/j.coph.2006.09.007

3 Hinkel, R. et al. Diabetes Mellitus-Induced Microvascular Destabilization in the Myocardium. J Am Coll Cardiol 69, 131–143 (2017). 10.1016/j.jacc.2016.10.058

4 Lang, R. A. & Bishop, J. M. Macrophages are required for cell death and tissue remodeling in the developing mouse eye. Cell 74, 453–462 (1993). 10.1016/0092-8674(93)80047-i

5 Lobov, I. B. et al. WNT7b mediates macrophage-induced programmed cell death in patterning of the vasculature. Nature 437, 417–421 (2005). 10.1038/nature03928

6 Reynolds, L. P., Killilea, S. D. & Redmer, D. A. Angiogenesis in the female reproductive system. FASEB J 6, 886–892 (1992).

7 Koller, G. M. et al. Proinflammatory Mediators, IL (Interleukin)-1beta, TNF (Tumor Necrosis Factor) alpha, and Thrombin Directly Induce Capillary Tube Regression. Arterioscler Thromb Vasc Biol 40, 365–377 (2020). 10.1161/ATVBAHA.119.313536

8 Control, C. f. D. Chronic Kidney Disease in the United States, 2023, <https://www.cdc.gov/kidney-disease/php/data-research/index.html> (2024).

9 Kida, Y. Peritubular Capillary Rarefaction: An Underappreciated Regulator of CKD Progression. Int J Mol Sci 21 (2020). 10.3390/ijms21218255

10 Kida, Y., Ieronimakis, N., Schrimpf, C., Reyes, M. & Duffield, J. S. EphrinB2 reverse signaling protects against capillary rarefaction and fibrosis after kidney injury. J Am Soc Nephrol 24, 559–572 (2013). 10.1681/ASN.2012080871

11 Babickova, J. et al. Regardless of etiology, progressive renal disease causes ultrastructural and functional alterations of peritubular capillaries. Kidney Int 91, 70–85 (2017). 10.1016/j.kint.2016.07.038

12 Gaupp, C. et al. Reconfiguration and loss of peritubular capillaries in chronic kidney disease. Sci Rep 13, 19660 (2023). 10.1038/s41598-023-46146-4

13 Li, S. et al. Proximal tubule PPARalpha attenuates renal fibrosis and inflammation caused by unilateral ureteral obstruction. Am J Physiol Renal Physiol 305, F618–627 (2013). 10.1152/ajprenal.00309.2013

14 Li, Z. I. et al. C-reactive protein promotes acute renal inflammation and fibrosis in unilateral ureteral obstructive nephropathy in mice. Lab Invest 91, 837–851 (2011). 10.1038/labinvest.2011.42

15 Chen, X. et al. Role of Reactive Oxygen Species in Tumor Necrosis Factor-alpha Induced Endothelial Dysfunction. Curr Hypertens Rev 4, 245–255 (2008). 10.2174/157340208786241336

16 Fajardo, L. F., Kwan, H. H., Kowalski, J., Prionas, S. D. & Allison, A. C. Dual role of tumor necrosis factor-alpha in angiogenesis. Am J Pathol 140, 539–544 (1992).

17 Karsan, A., Yee, E. & Harlan, J. M. Endothelial cell death induced by tumor necrosis factor-alpha is inhibited by the Bcl-2 family member, A1. J Biol Chem 271, 27201–27204 (1996). 10.1074/jbc.271.44.27201

18 Mountain, D. J., Singh, M. & Singh, K. Interleukin-1beta-mediated inhibition of the processes of angiogenesis in cardiac microvascular endothelial cells. Life Sci 82, 1224–1230 (2008). 10.1016/j.lfs.2008.04.008

19 Mukohda, M. et al. Endothelial PPAR-gamma provides vascular protection from IL-1beta-induced oxidative stress. Am J Physiol Heart Circ Physiol 310, H39–48 (2016). 10.1152/ajpheart.00490.2015

20 Pohlman, T. H. & Harlan, J. M. Human endothelial cell response to lipopolysaccharide, interleukin-1, and tumor necrosis factor is regulated by protein synthesis. Cell Immunol 119, 41–52 (1989). 10.1016/0008-8749(89)90222-0

21 Schafer, C. M. et al. Cytokine-Mediated Degradation of the Transcription Factor ERG Impacts the Pulmonary Vascular Response to Systemic Inflammatory Challenge. Arterioscler Thromb Vasc Biol 43, 1412–1428 (2023). 10.1161/ATVBAHA.123.318926

22 Wang, Y. et al. Ephrin-B2 controls VEGF-induced angiogenesis and lymphangiogenesis. Nature 465, 483–486 (2010). 10.1038/nature09002

23 Chevalier, R. L., Forbes, M. S. & Thornhill, B. A. Ureteral obstruction as a model of renal interstitial fibrosis and obstructive nephropathy. Kidney Int 75, 1145–1152 (2009). 10.1038/ki.2009.86

24 Hesketh, E. E. et al. A murine model of irreversible and reversible unilateral ureteric obstruction. J Vis Exp (2014). 10.3791/52559

25 Lopez De Padilla, C. M., et al. Picrosirius Red Staining: Revisiting Its Application to the Qualitative and Quantitative Assessment of Collagen Type I and Type III in Tendon. J Histochem Cytochem 69, 633–643 (2021). 10.1369/00221554211046777

26 Whittaker, P., Kloner, R. A., Boughner, D. R. & Pickering, J. G. Quantitative assessment of myocardial collagen with picrosirius red staining and circularly polarized light. Basic Res Cardiol 89, 397–410 (1994). 10.1007/BF00788278

27 Choi, Y. J. et al. Peritubular capillary loss is associated with chronic tubulointerstitial injury in human kidney: altered expression of vascular endothelial growth factor. Hum Pathol 31, 1491–1497 (2000). 10.1053/hupa.2000.20373

28 Han, W. K., Bailly, V., Abichandani, R., Thadhani, R. & Bonventre, J. V. Kidney Injury Molecule-1 (KIM-1): a novel biomarker for human renal proximal tubule injury. Kidney Int 62, 237–244 (2002). 10.1046/j.1523-1755.2002.00433.x

29 Zhang, P. L. & Liu, M. L. From acute tubular injury to tubular repair and chronic kidney diseases - KIM-1 as a promising biomarker for predicting renal tubular pathology. Curr Res Physiol 8, 100152 (2025). 10.1016/j.crphys.2025.100152

30 Lin, P. K. & Davis, G. E. Extracellular Matrix Remodeling in Vascular Disease: Defining Its Regulators and Pathological Influence. Arterioscler Thromb Vasc Biol 43, 1599–1616 (2023). 10.1161/ATVBAHA.123.318237

31 Huang, R., Fu, P. & Ma, L. Kidney fibrosis: from mechanisms to therapeutic medicines. Signal Transduct Target Ther 8, 129 (2023). 10.1038/s41392-023-01379-7

32 Nan, Q. Y. et al. Pathogenesis and management of renal fibrosis induced by unilateral ureteral obstruction. Kidney Res Clin Pract 43, 586–599 (2024). 10.23876/j.krcp.23.156

33 Panizo, S. et al. Fibrosis in Chronic Kidney Disease: Pathogenesis and Consequences. Int J Mol Sci 22 (2021). 10.3390/ijms22010408

34 LeBleu, V. S. et al. Origin and function of myofibroblasts in kidney fibrosis. Nat Med 19, 1047–1053 (2013). 10.1038/nm.3218

35 Matsumoto, M. et al. Hypoperfusion of peritubular capillaries induces chronic hypoxia before progression of tubulointerstitial injury in a progressive model of rat glomerulonephritis. J Am Soc Nephrol 15, 1574–1581 (2004). 10.1097/01.asn.0000128047.13396.48

36 Mayer, G. Capillary rarefaction, hypoxia, VEGF and angiogenesis in chronic renal disease. Nephrol Dial Transplant 26, 1132–1137 (2011). 10.1093/ndt/gfq832

37 Tanaka, S., Tanaka, T. & Nangaku, M. Hypoxia as a key player in the AKI-to-CKD transition. Am J Physiol Renal Physiol 307, F1187–1195 (2014). 10.1152/ajprenal.00425.2014

38 Chang, F. C. et al. Angiopoietin-2 inhibition attenuates kidney fibrosis by hindering chemokine C-C motif ligand 2 expression and apoptosis of endothelial cells. Kidney Int 102, 780–797 (2022). 10.1016/j.kint.2022.06.026

39 Kim, W. et al. COMP-angiopoietin-1 ameliorates renal fibrosis in a unilateral ureteral obstruction model. J Am Soc Nephrol 17, 2474–2483 (2006). 10.1681/ASN.2006020109

40 Singh, S. et al. Tubular Overexpression of Angiopoietin-1 Attenuates Renal Fibrosis. PLoS One 11, e0158908 (2016). 10.1371/journal.pone.0158908

41 Lin, S. L. et al. Targeting endothelium-pericyte cross talk by inhibiting VEGF receptor signaling attenuates kidney microvascular rarefaction and fibrosis. Am J Pathol 178, 911–923 (2011). 10.1016/j.ajpath.2010.10.012

42 Sainson, R. C. et al. TNF primes endothelial cells for angiogenic sprouting by inducing a tip cell phenotype. Blood 111, 4997–5007 (2008). 10.1182/blood-2007-08-108597

43 Frater-Schroder, M., Risau, W., Hallmann, R., Gautschi, P. & Bohlen, P. Tumor necrosis factor type alpha, a potent inhibitor of endothelial cell growth in vitro, is angiogenic in vivo. Proc Natl Acad Sci U S A 84, 5277–5281 (1987). 10.1073/pnas.84.15.5277

44 Kowluru, R. A. & Odenbach, S. Role of interleukin-1beta in the development of retinopathy in rats: effect of antioxidants. Invest Ophthalmol Vis Sci 45, 4161–4166 (2004). 10.1167/iovs.04-0633

45 Voronov, E. et al. IL-1 is required for tumor invasiveness and angiogenesis. Proc Natl Acad Sci U S A 100, 2645–2650 (2003). 10.1073/pnas.0437939100

46 Martinez-Klimova, E., Aparicio-Trejo, O. E., Tapia, E. & Pedraza-Chaverri, J. Unilateral Ureteral Obstruction as a Model to Investigate Fibrosis-Attenuating Treatments. Biomolecules 9 (2019). 10.3390/biom9040141

47 Loganathan, K. et al. Angiopoietin-1 deficiency increases renal capillary rarefaction and tubulointerstitial fibrosis in mice. PLoS One 13, e0189433 (2018). 10.1371/journal.pone.0189433

48 Ren, Z. et al. Contribution of alterations in peritubular capillary density and microcirculation to the progression of tubular injury and kidney fibrosis. J Pathol 266, 95–108 (2025). 10.1002/path.6414

49 Chen, X. et al. Overexpression of Heme Oxygenase-1 Prevents Renal Interstitial Inflammation and Fibrosis Induced by Unilateral Ureter Obstruction. PLoS One 11, e0147084 (2016). 10.1371/journal.pone.0147084

50 Lin, S. L., Kisseleva, T., Brenner, D. A. & Duffield, J. S. Pericytes and perivascular fibroblasts are the primary source of collagen-producing cells in obstructive fibrosis of the kidney. Am J Pathol 173, 1617–1627 (2008). 10.2353/ajpath.2008.080433

51 Xavier, S. et al. Curtailing endothelial TGF-beta signaling is sufficient to reduce endothelial-mesenchymal transition and fibrosis in CKD. J Am Soc Nephrol 26, 817–829 (2015). 10.1681/ASN.2013101137

52 Maleszewska, M. et al. IL-1beta and TGFbeta2 synergistically induce endothelial to mesenchymal transition in an NFkappaB-dependent manner. Immunobiology 218, 443–454 (2013). 10.1016/j.imbio.2012.05.026

53 Yoshimatsu, Y. et al. TNF-alpha enhances TGF-beta-induced endothelial-to-mesenchymal transition via TGF-beta signal augmentation. Cancer Sci 111, 2385–2399 (2020). 10.1111/cas.14455

54 Schafer, C. M. et al. An inhibitor of endothelial ETS transcription factors promotes physiologic and therapeutic vessel regression. Proc Natl Acad Sci U S A 117, 26494–26502 (2020). 10.1073/pnas.2015980117

55 Ma, E. et al. Targeting endothelial ERG to mitigate vascular regression in retinopathies. Proc Natl Acad Sci U S A 122, e2507194122 (2025). 10.1073/pnas.2507194122

56 Paavonsalo, S., Hariharan, S., Lackman, M. H. & Karaman, S. Capillary Rarefaction in Obesity and Metabolic Diseases-Organ-Specificity and Possible Mechanisms. Cells 9 (2020). 10.3390/cells9122683

57 Miranda, M., Balarini, M., Caixeta, D. & Bouskela, E. Microcirculatory dysfunction in sepsis: pathophysiology, clinical monitoring, and potential therapies. Am J Physiol Heart Circ Physiol 311, H24–35 (2016). 10.1152/ajpheart.00034.2016

58 Zlokovic, B. V. Neurovascular pathways to neurodegeneration in Alzheimer’s disease and other disorders. Nat Rev Neurosci 12, 723–738 (2011). 10.1038/nrn3114

59 Mohammed, S. F. et al. Coronary microvascular rarefaction and myocardial fibrosis in heart failure with preserved ejection fraction. Circulation 131, 550–559 (2015). 10.1161/CIRCULATIONAHA.114.009625

60 Osiaevi, I. et al. Persistent capillary rarefication in long COVID syndrome. Angiogenesis 26, 53–61 (2023). 10.1007/s10456-022-09850-9

61 Saifi, M. A., Annaldas, S. & Godugu, C. A direct thrombin inhibitor, dabigatran etexilate protects from renal fibrosis by inhibiting protease activated receptor-1. Eur J Pharmacol 893, 173838 (2021). 10.1016/j.ejphar.2020.173838

62 Erreger, K. et al. Role of protease-activated receptor 4 in mouse models of acute and chronic kidney injury. Am J Physiol Renal Physiol 326, F219–F226 (2024). 10.1152/ajprenal.00162.2023

