## Supplemental Figs and Tables for "TNFα and Endothelial IL-1 Receptor Signaling Drive Peritubular Capillary Regression and Fibrosis in Obstructive Kidney Injury"

**This PDF file includes:**

Figures S1 to S5

Tables S1 to S3

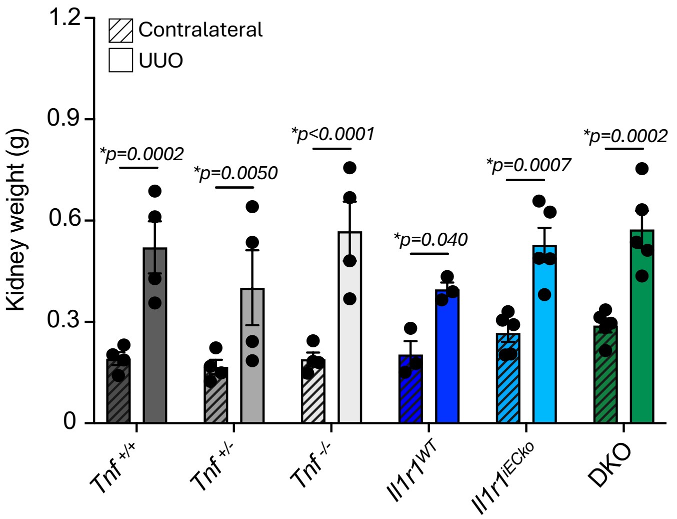


**Figure S1: Kidney weight is increased after UUO in all genotypes.**
Kidney weights of contralateral and UUO kidneys were measured in all genotypes. Each dot represents an individual animal; N=3-5. Data are presented as mean ± SEM. Data were analyzed using two-way ANOVA and post hoc comparisons.

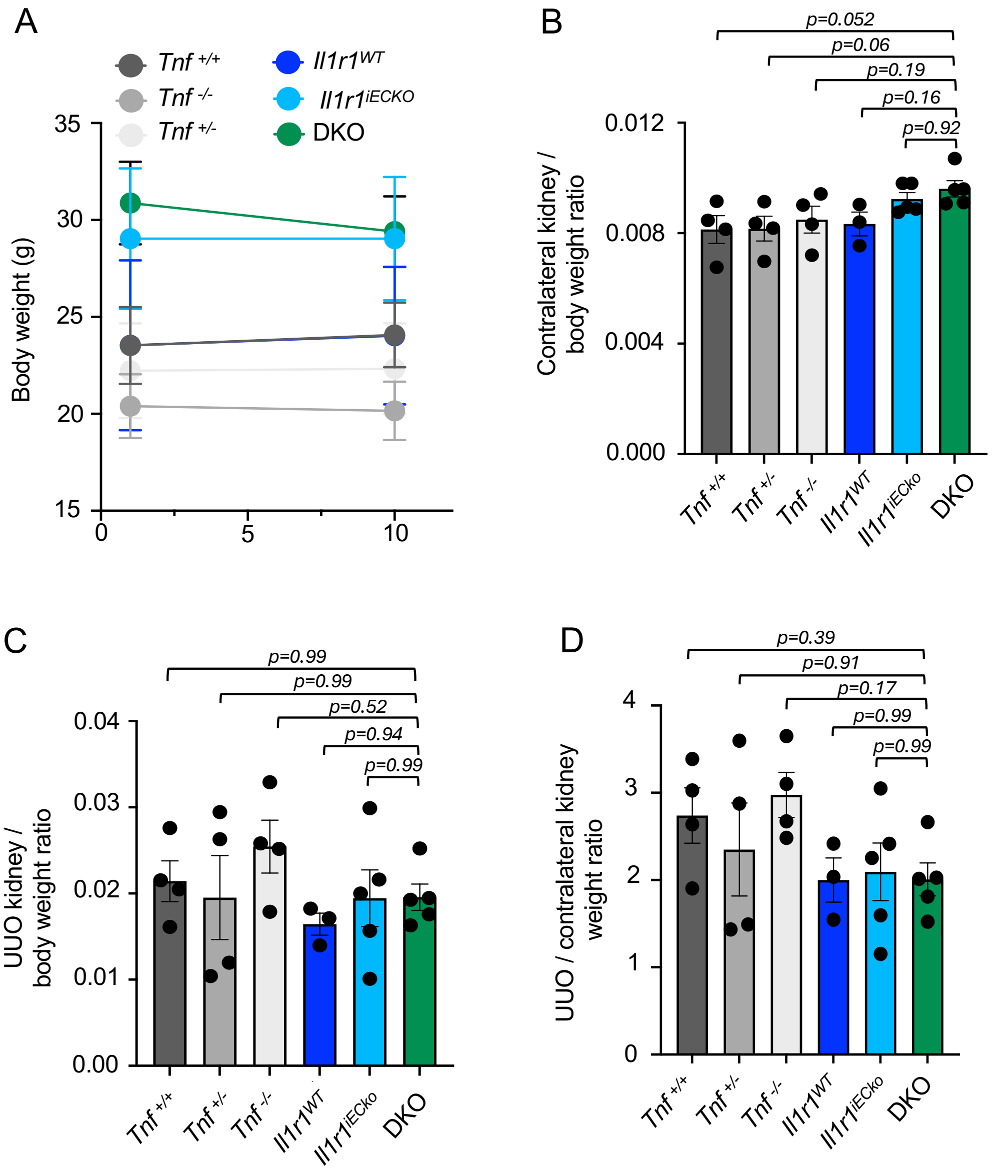


**Figure S2: TNFα and endothelial IL-1R1 signaling does not influence UUO-induced kidney and body weight changes.**

(A) Body weight measurements at days 0 and 10 post-UUO across all genotypes. N=3-5. (B) Contralateral kidney-to-body weight ratios, (C) UUO kidney-to-body weight ratios and (D) UUO/contralateral kidney weight ratios across all genotypes. For B-D, each dot represents one mouse; N=3-5. Data are presented as mean ± SEM. Data were analyzed using one-way ANOVA and post hoc comparisons.

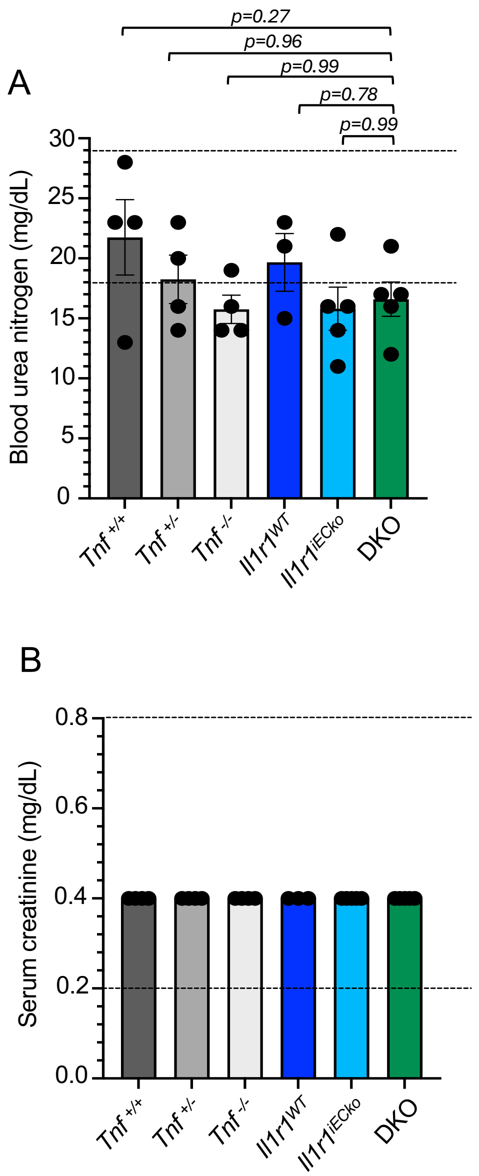


**Figure S3. Markers of kidney function remain stable following UUO across genotypes.**

(A) Blood urea nitrogen (BUN) levels measured in serum in mice across all genotypes. Horizontal dashed lines indicate the normal reference range (18–29 mg/dL). (B) Serum creatinine levels were measured across all genotypes, with values remaining within the normal range (dashed lines; 0.2–0.8 mg/dL). Each data point represents an individual animal; N=3-5. Data are presented as mean ± SEM. Data were analyzed using one-way ANOVA and post hoc comparisons.

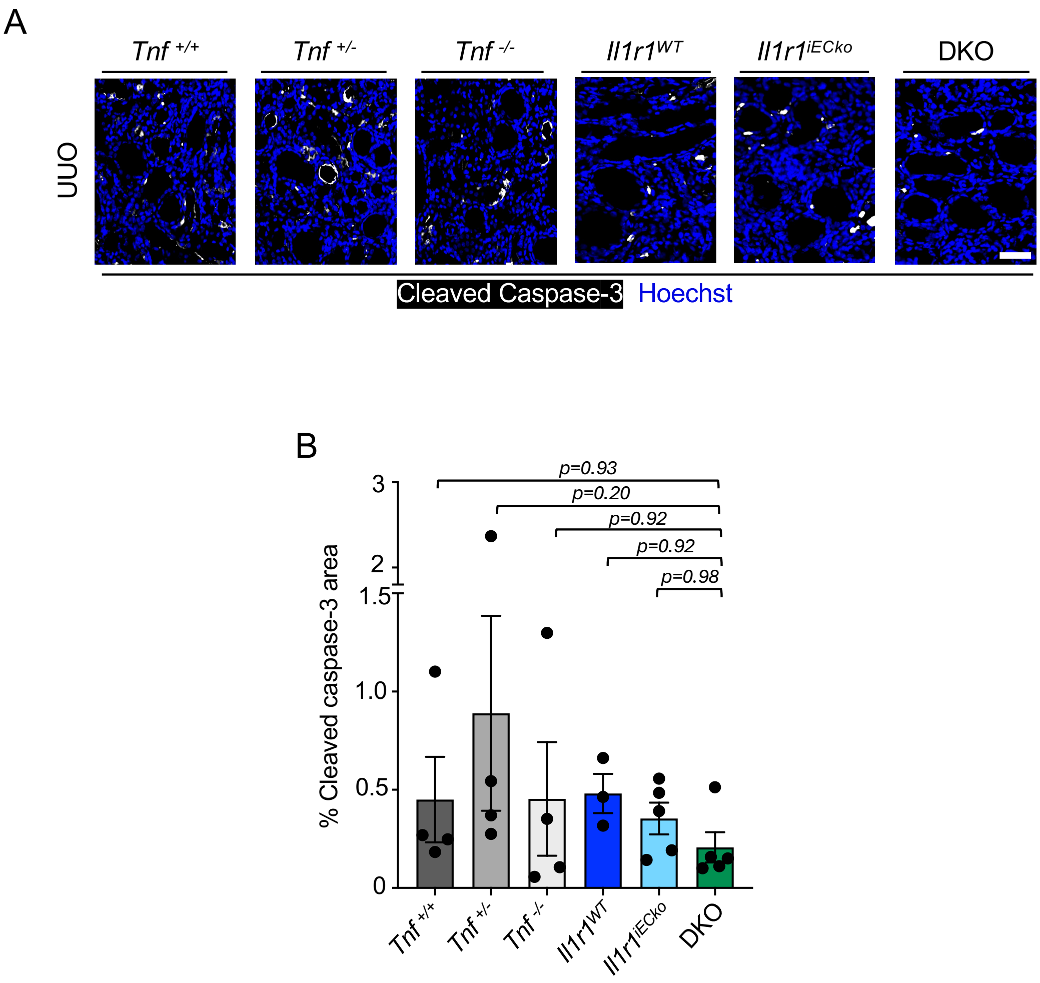


**Figure S4. Cell death is not significantly affected across genotypes following UUO.** (A) Representative immunofluorescence images of UUO kidney sections stained for cleaved caspase-3 (white) and Hoechst (blue) in the indicated genotypes. Scale bar = 20 µm. (B) Quantification of cleaved caspase-3–positive area as a percentage of total area. Data represent mean ± SEM. Each data point represents an individual animal; N=3-5. 3-4 images were quantified per animal. Data were analyzed using one-way ANOVA.

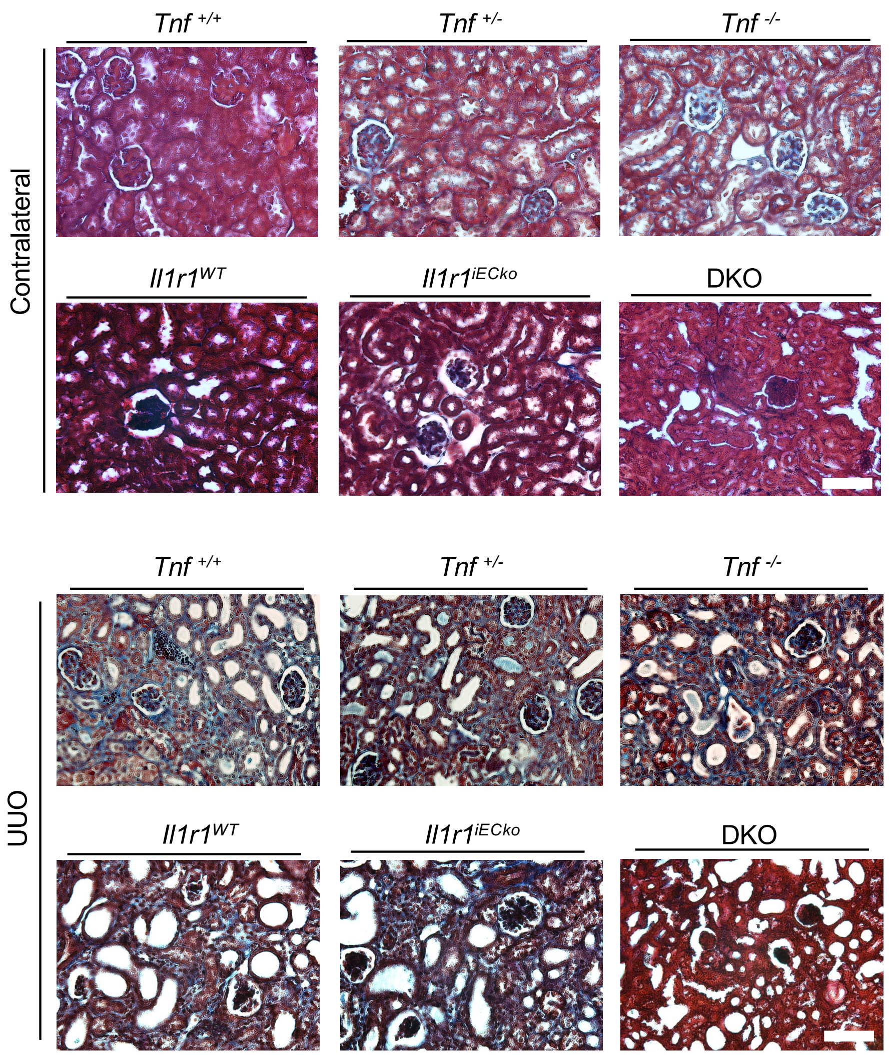


**Figure S5. Histological assessment of kidney fibrosis.**
Masson's trichrome staining of contralateral and UUO kidneys with collagen fibers depicted in blue. Scale bar = 50 μm.

**Table S1: List of primers used for genotyping**

| Primer Name | Sequence | Allele |
|---|---|---|
| *Il1r1-flox-F* | 5'-GAAAAGTGCTAGAACATCCTTTGAG | *Il1r1-flox* |
| *Il1r1-flox-R* | 5'-GTACCAATGGAGGCCAGAAG | |
| *Tnf-WT-F* | 5`-TAGCCAGGAGGGAGAACAGA | *Tnf-KO* |
| *Tnf-WT-R* | 5`-AGTGCCTCTTCTGCCAGTTC | |
| *Tnf-MUT-R* | 5`-CGTTGGCTACCCGTGATATT | |
| *iCdh5-Cre-F* | 5`-GTACCAATGGAGGCCAGAAG | *iCdh5-Cre$^{ERT2}$* |
| *iCdh5-Cre-R* | 5’-CGAACCTGGTCGAAATCAGT | |
| Control-F | 5’-CGAACCTGGTCGAAATCAGT | |
| Control-R | 5’-GTAGGTGGAAATTCTAGCATCATCC | |

**Table S2: Major Resources Table**

### Animals (in vivo studies)

| Species | Vendor or Source | Background Strain | Sex | Persistent ID / URL |
|---|---|---|---|---|
| Mouse | Taconic | *Cdh5(PAC)-Cre*$^{ERT2}$ | M/F | #13073 |
| Mouse | Jackson Laboratory | *Il1r1*$^{flox}$ | M/F | #028398 |
| Mouse | Jackson Laboratory | *Tnf* $^{-/-}$ | M/F | #003008 |

### Primary Antibodies

| Target antigen | Vendor or Source | Catalog # | Working concentration | Persistent ID / URL |
|---|---|---|---|---|
| CD31 | R&D Systems | AF3628 | 1:100 | RRID:AB_2161028 |
| aSMA | Sigma | C6198 | 1:250 | RRID:AB_476856 |
| KIM-1 | R&D Systems | AF1817 | 1:100 | RRID:AB_2116446 |
| Cleaved caspase-3 | Cell Signaling | 9661 | 1:400 | RRID:AB_2341188 |

### Secondary Antibodies

| Host and Target species | Fluorophore or conjugate | Vendor or Source | Catalog # | Working concentration | Persistent ID / URL |
|---|---|---|---|---|---|
| Donkey anti-Rabbit | Cy5 | Jackson ImmunoResearch | 711-175-152 | 1:500 | RRID:AB_2340607 |
| Donkey anti-Rat | Alexa488 | Thermo Fisher | A-21208 | 1:500 | RRID:AB_2535794 |
| Donkey anti-Goat | Cy3 | Jackson ImmunoResearch | 705-165-003 | 1:500 | RRID:AB_2340411 |

**Table S3: Experimental mouse information**

**Figure 1D, 2C, 3B, 4B, S1, S2A-D, S3A-B, S4B**

| Groups | Sex | Age | Number (prior to experiment) (M=male, F=female) | Number (after termination) (M=male, F=female) | Littermates (Yes/No) |
|---|---|---|---|---|---|
| *Tnf* $^{+/+}$ | M/F | 12 weeks old | 2M, 2F | 2M, 2F | Yes |
| *Tnf* $^{+/-}$ | M/F | 12 weeks old | 1M, 3F | 1M, 3F | Yes |
| *Tnf* $^{-/-}$ | M/F | 12 weeks old | 2M, 2F | 2M, 2F | Yes |
| *Il1r1*$^{WT}$ | M/F | 12 weeks old | 1M, 2F | 1M, 2F | Yes |
| *Il1r1*$^{iECko}$ | M/F | 12 weeks old | 3M, 2F | 3M, 2F | Yes |
| *Tnf* $^{-/-}$; *Il1r1*$^{iECKO}$ (DKO) | M/F | 12 weeks old | 3M, 2F | 3M, 2F | Yes |
